# Patterns of unwanted biological and technical expression variation across 49 human tissues

**DOI:** 10.1101/2023.03.09.531935

**Authors:** Tim O. Nieuwenhuis, Hunter H. Giles, Matthew N. McCall, Marc K. Halushka

## Abstract

All tissue-based gene expression studies are impacted by biological and technical sources of variation. Numerous methods are used to normalize and batch correct these datasets. A more accurate understanding of all causes of variation could further optimize these approaches. We used 17,282 samples from 49 tissues in the Genotype Tissue Expression (GTEx) dataset (v8) to investigate patterns and causes of expression variation. Transcript expression was normalized to Z-scores and only the most variable 2% of transcripts were evaluated and clustered based on co-expression patterns. Clustered gene sets were solved to different biological or technical causes related to metadata elements and histologic images. We identified 522 variable transcript clusters (median 11 per tissue) across the samples. Of these, 64% were confidently explained, 15% were likely explained, 7% were low confidence explanations and 14% had no clear cause. Common causes included sex, sequencing contamination, immunoglobulin diversity, and compositional tissue differences. Less common biological causes included death interval (Hardy score), muscle atrophy, diabetes status, and menopause. Technical causes included brain pH and harvesting differences. Many of the causes of variation in bulk tissue expression were identifiable in the Tabula Sapiens dataset of single cell expression. This is the largest exploration of the underlying sources of tissue expression variation. It uncovered expected and unexpected causes of variable gene expression. These identified sources of variation will inform which metadata to acquire with tissue harvesting and can be used to improve normalization, batch correction, and analysis of both bulk and single cell RNA-seq data.

## INTRODUCTION

Technical and biological variability result in expression heterogeneity across tissue samples, impacting the analysis of large expression datasets. Identifying and correcting for unwanted variation remains one of the great challenges of –omic style studies. Human tissue studies, whether single-cell, single-nucleus, or bulk RNA-seq are all confounded by a large number of factors related to procurement, experimental steps, and life experiences of the tissue donors. As seen repeatedly, how this variation is accounted for will deeply affect the expression estimates in cells and tissues and impact the true positive discovery of differentially expressed genes (Leek et al. 2012; Somekh et al. 2019; Chen et al. 2020).

Strategies to correct for unwanted variation fall into two general categories – inferred batch and known batch correction, and often times these approaches are combined. Inferred batch methods such as surrogate variable analysis (SVA), Seurat, and PEER factors are generally agnostic to the cause of sample heterogeneity and create covariates that regress out/correct for variation without regard to the reason for heterogeneity (Leek et al. 2012; Stegle et al. 2012; Butler et al. 2018). These inferred variables tend to remove both biological and technical factors, unless the biological difference of interest is included in the model. Alternatively, with batch correction methods, such as ComBat, and linear model based differential expression methods, such as edgeR, limma, and DESeq2, one can make a direct adjustment for known sources of unwanted variation only (Johnson et al. 2007; Robinson et al. 2010; Love et al. 2014; Ritchie et al. 2015). However, these methods are limited to the technical and biological variables that are recorded, missing possible confounding factors. Generally, these batch correction methods correct for known technical factors while leaving biological drivers of variation uncorrected.

A direct adjustment approach, when experimental metadata is well-captured and incorporated into a batch correct strategy, is consistent with the general assumption that technical variability should be removed. Conversely, less has been settled on how to approach variable biological factors, other than identifying and not correcting for, the biological variable of interest. Well-known technical factors include RNA quality, sequencing dates, and reagent differences. Confounding biological variables are poorly characterized. Commonly known and corrected variables include age and sex, but some variables, including the manner of death and medication usage, may impact gene expression and might additionally be attractive for correction approaches. In the absence of knowing what variables to capture and correct for, our bulk and single cell RNA-seq studies will need surrogate variable approaches which undoubtedly alter the interpretation of the sequencing projects.

The Genotype Tissue Expression (GTEx) project is a useful resource to understand technical and biologic causes of expression heterogeneity (McCall et al. 2016; Donovan et al. 2020; Nieuwenhuis et al. 2020). Although originally designed to identify expression Quantitative Trait Loci (eQTLs), many investigators have found GTEx to be a powerful tool for secondary analyses (Somekh et al. 2019; Donovan et al. 2020; Nieuwenhuis et al. 2020; Luca et al. 2021; Nieuwenhuis et al. 2021). GTEx contains bulk RNA-seq data from up to 948 donors across 54 tissues. GTEx augments these data with digitally scanned tissue images, fairly rich donor clinical history, and robust technical data points related to tissue procurement, processing, and sequencing (Consortium et al. 2017). We previously used an earlier version of the GTEx database to investigate drivers of variation in lung tissue (McCall et al. 2016) identifying procurement location (medial vs lateral) and ventilator use at the time of death as two unexpected major drivers of variability. GTEx has also been used to demonstrate the advantage of a linear regression model approach using known confounders versus approaches that estimate hidden confounders (Somekh et al. 2019). Therefore, understanding the biological and technical drivers of tissue heterogeneity, so that they can be employed in batch correction, is of great value.

In this project we utilize the entire GTEx collection (v8) to identify common and unique patterns of variation across the majority of the tissues. This work uncovered unexpected biological, phenotypic and technical causes of expression variation. Many of the drivers of variation that were uncovered are not generally captured metadata points in single cell and bulk studies, but should be collected, in light of the discoveries made herein.

## RESULTS

### Establishment of co-variable gene clusters

In total, 49 tissues across 17,282 samples (minimum [min] = 85, kidney cortex; maximum [max] = 803, skeletal muscle; median 258) were analyzed (**Supplemental Table S1**). Five tissues (bladder, endocervix, ectocervix, fallopian tube, and kidney medulla) were dropped due to a lack of samples. After variance stabilizing transformation (VST) of transcript (e.g. gene, pseudogene, lncRNA) expression for each sample, an intra-tissue minimum expression filter (mean VST count ≥5 across samples), passed a median of 18,002 transcripts per tissue (min = 13,954, whole blood; max = 25,108, testis). From each tissue transcript set, a variance filter captured the top 2% most variable transcripts in each tissue. Passing transcripts had a median gene variance of 2.36 across tissues (min = 0.99 cerebellum, max = 8.53 transverse colon) identifying a median of 361 transcripts per tissue (total = 4,961 transcripts, min = 280, whole blood; max = 503, testis).

All variable transcripts underwent hierarchical and agglomerative clustering, as described in the methods, and in some cases additional manual curation, into 522 clusters (cluster min size = 6 transcripts) (**Fig. 1A-B**; **Supplemental Table S2**, **Supplemental Figs. S1-49**). Some variable transcripts did not correlate with other transcripts after clustering (min = 6, Colon – transverse; max = 159, Brain – cerebellar hemisphere) ( **Supplemental Table S3**). Clusters were arbitrarily named using the GTEx tissue naming convention and sequential alphabetical numbers with characters “.1” and “.2” added to clusters when transcripts are both positively and negatively correlated (e.g. ADPSBQ.J.1, ADPSBQ.J.2, KDNCTX.A) (Nieuwenhuis et al. 2021). As in our earlier work, for each transcript we transformed the normalized and variance stabilized gene expression values to z-scores by subtracting the mean and then dividing by the median absolute deviation. To produce sample-specific estimates of average expression for a cluster, these z-scores were averaged across all genes in the cluster (McCall et al. 2016). A median of 11 unique clusters, passing these thresholds, were present in each tissue (min. = 4, testis/vagina; max. = 22, uterus), with the etiology of each cluster being generally independently correlated with a technical or biological factor. Of the 522 variable transcript clusters, 64% were confidently solved, 15% were likely solved and 7% had low confidence in a cause and 14% had no clear etiology.

**Figure 1.**
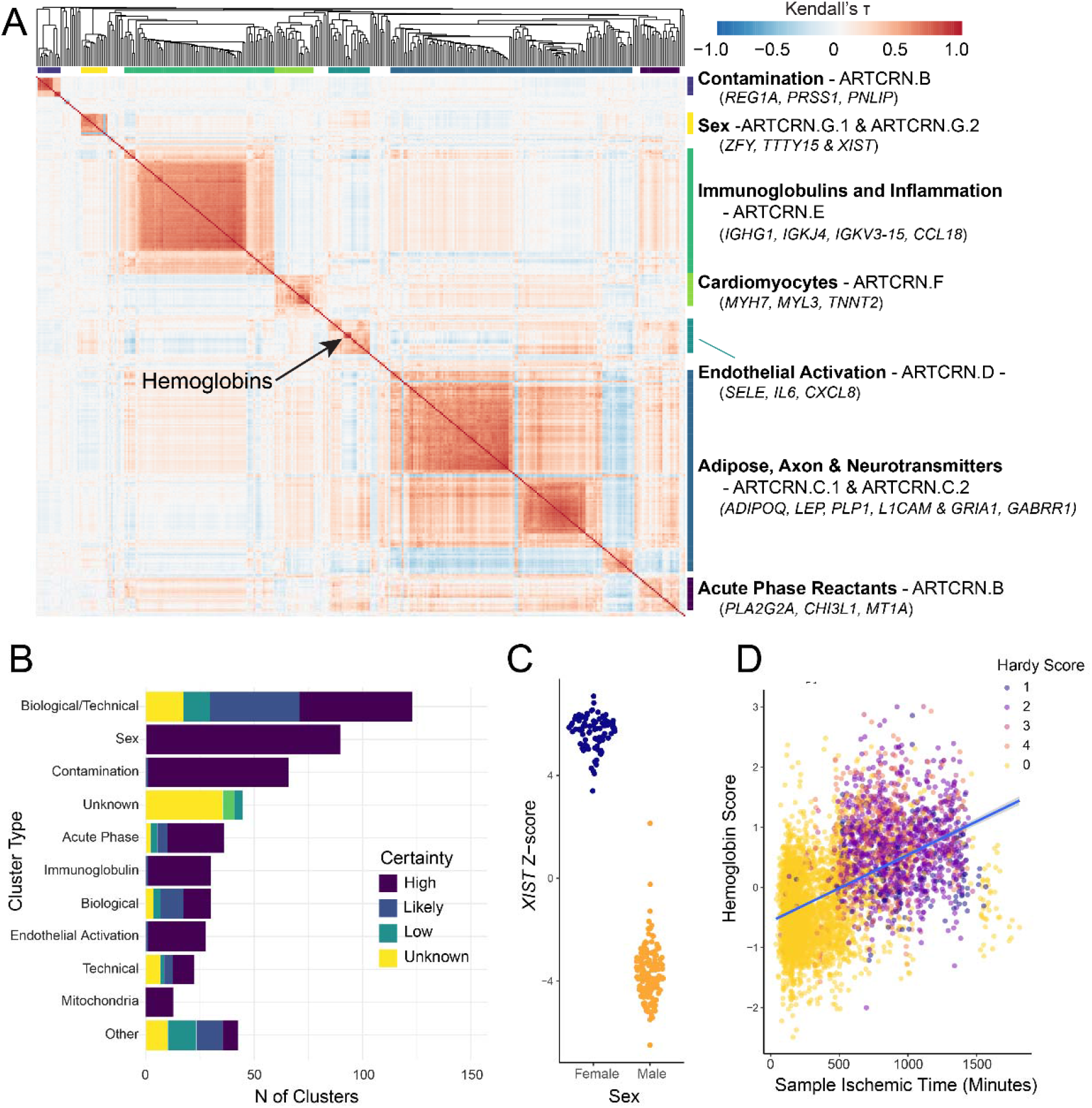
Global trends in bulk RNA-seq tissues from GTEx. (**A)** A representative heatmap from the coronary artery (ARTCRN;N = 240 samples) of a Kendall’s τ correlation matrix of high variance transcripts (N = 355 high variance transcripts) identifies 9 clusters (7 clusters with 2 having positive and negative subclusters). Red indicates positive correlation and blue indicates negative correlations. Above the heatmap is the dendrogram clustering the high variance transcripts on 1 - |Kendall’s τ Correlation|. Correlates/causes of clusters and representative genes are provided. **(B)** Cluster type and confidence of all 522 clusters across 49 tissues. **(C)**A sina plot of expression Z-score distribution of *XIST*, in males and females (N = 146 and 94 respectively). **(D)** Expression of a collapsed mean Z-score of all hemoglobin genes ( *HBB, HBA1, HBA2,*and *HBD*) vs ischemic time for 12 tissues (Ferreira et al. 2018). The samples are colored by Hardy Score (death interval).

### Frequent, expected transcript clusters across multiple tissues

Several causes of transcript clusters being variable across tissues were noted. Some of these were relatively straightforward to identify. The most common associations of these clusters were to subject sex, immunoglobulins, and sequencing contamination. A sex cluster comprised of Y chromosome genes (*TTTY15*, *ZFY*, *RPS4Y1*, etc.) and inversely correlated with the X-inactivation transcript *XIST*, clearly distinguished male and female subjects across 43 of the tissues (**Fig. 1C**).

A variable immunoglobulin expression cluster was noted across 30 tissues. In total, 136 immunoglobulin genes were included in clusters. These ranged from constant structural elements (e.g. *IGHA1*, *IGHG1*, *IGKJ*)*1*to specific variable regions (e.g. *IGHV7-4-1*, *IGHV3-23*). These collections reflect both an inconstant number of plasma cells within tissues and individual differences in immunoglobulin variable regions (Mandric et al. 2020).

The hemoglobin genes (*HBA1*, *HBA2*, *HBB*, and *HBD*) were each identified as variable in 30-43 tissues, depending on the specific gene. As our minimum threshold to form a cluster is 5 transcripts, these genes were often unclustered ( **Supplemental Tables S2, S3**), but when in a named cluster (N=19), they were associated with non-red blood cell specific transcripts. An increase in *HBB*transcript levels has been previously noted to associate with ischemic time (Ferreira et al. 2018). That correlation is also true for all hemoglobin genes in a linear mixed model while adjusting for multiple covariates including Hardy Score (**Fig. 1D; Supplemental Fig. S50;** *HBB*, *HBA1*, *HBA2*, *HBD*; respective ischemic time p-value: 4.6e-4, 6.2e-4, 9.9e-05, 0.0069). Hardy score (range 0-4), is a variable that bins terminal phases of death between acute (<10 minutes, score 1), prolonged (>1 day, score 4) or on a ventilator at death (score 0). In general, hemoglobin variability is most associated with the consistency of washing tissues, with less washing resulting in more adherent blood. Secondarily, an increase with ischemic time likely reflects a mechanism in which endothelial cell integrity is reduced and more red blood cells simply become trapped in the tissue.

Sequencing contamination was also a common etiology of clusters. We previously demonstrated that library preparation-associated contamination can allow abundant genes from one GTEx tissue to be variability identified at low expression levels in a second GTEx tissue (Nieuwenhuis et al. 2020). This was primarily demonstrated with highly-abundant pancreas genes *PNLIP*, *PRSS1,*and *CELA3A*. We observed this contamination cluster in 38 tissues. We also identified a second contamination cluster in 25 tissues that contained genes including *ACTA1*, *MYL2,SFTPB*, and *KRT1*. Altogether, these frequent biological and technical drivers of variation accounted for 187 of the 522 clusters (36%) (**Supplemental Table S2**).

### Frequent, unexpected transcript clusters among multiple tissues

#### Hardy Score correlations

Beyond sex, immunoglobulins, and contamination, there were two common variable transcript clusters that correlated with Hardy Score, the measure of death interval. One frequent cluster contained tissue-based acute phase reactants including *PLA2G2A, CHI3L1, SAA1,* and *HP*(Birts et al. 2008; Abouelasrar Salama et al. 2020; Zhao et al. 2020; Gulhar et al. 2022) (**Fig. 2A**)*. PLA2G2A*, found variable in 28 tissues and in 25 clusters, was the main marker gene for this acute phase reactants cluster. Across the 25 (non-brain) clusters, there was a strong positive correlation between higher levels of the cluster genes and increasing death interval from acute death to hospitalized death (**Fig. 2B**).

**Figure 2.**
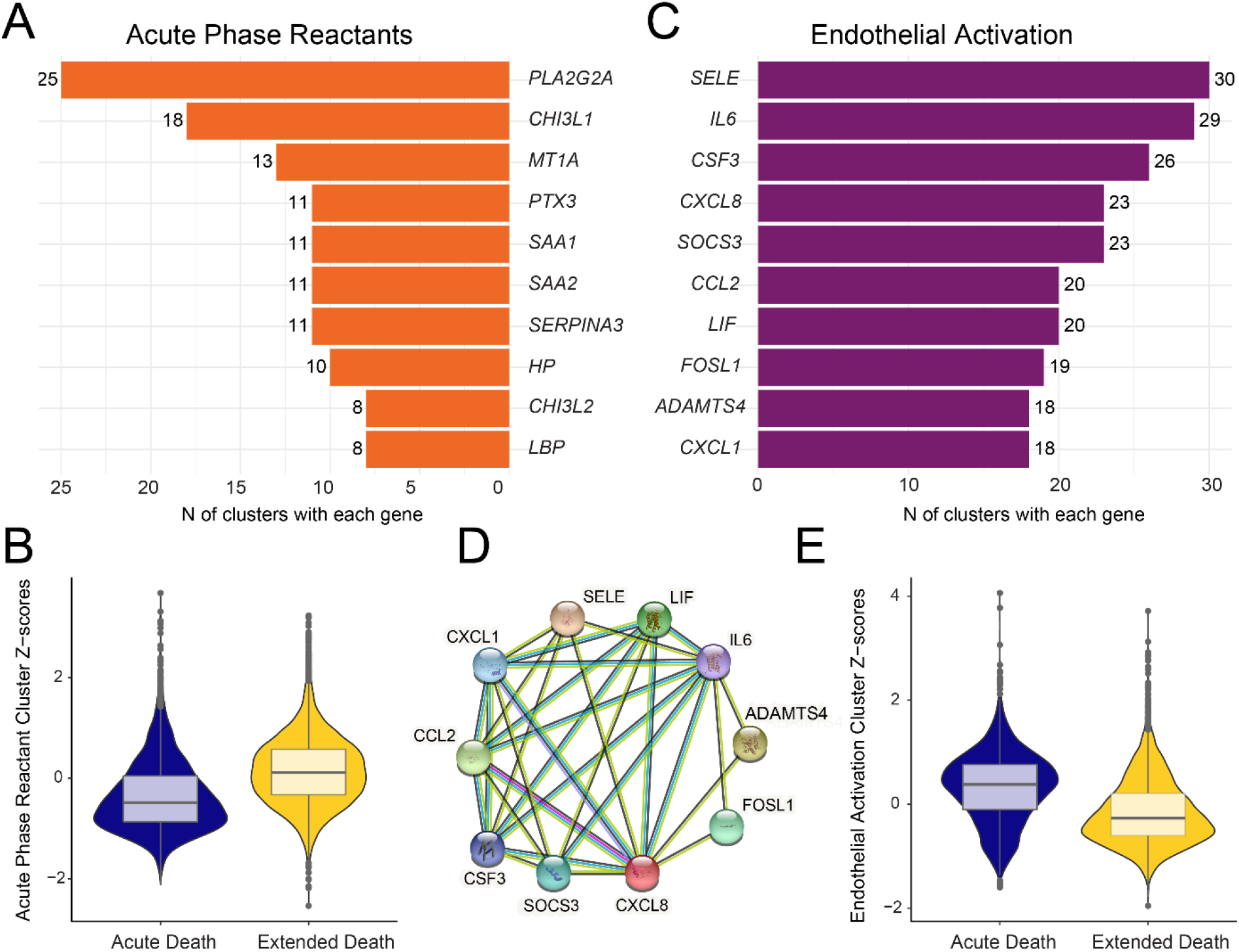
Acute phase reactant and endothelial activation clusters inversely correlate with Hardy Score. **(A)** The top 10 most frequent genes in the acute phase reactant cluster. **(B)** A violin plot/ boxplot of acute phase reactant cluster average Z-scores by Acute Death (Hardy = 1 or 2) or Extended Death (Hardy = 4 or 0). Brain clusters excluded. **(C)**The top 10 most frequent genes of the endothelial activation cluster. **(D)** A protein interaction map of the 10 endothelial activation genes (StringDB v11.5, protein-protein enrichment p-values: < 1.0e-16). **(E)** A violin plot/boxplot for endothelial activation cluster average Z-scores showing the opposite correlation to Hardy scores.

A second frequent cluster of transcripts related to Hardy score consistently contained *IL6* and *SELE.*This gene set is thought to be associated with endothelial activation and was notable for strong functional linkages across the genes (Chi et al. 2001; Su et al. 2012) ( **Fig. 2C, D**). Interestingly, there was a significant inverse correlation to Hardy score across non-brain tissues, where the highest cluster Z-scores correlated with the shortest death interval (**Fig. 2E**). Conversely, among the 9 brain tissues, there was a positive correlation of the highest Z-scores with the longest death intervals.

### Organ specific clusters

Numerous clusters were present with a set of genes/transcripts that were enriched for a particular organ or a shared set of organs. We identified biologic or technical reasons for 175 of these clusters. Many of these clusters indicated the heterogeneity of a particular cell type in the tissue, a type of sampling variability that can be the result of technical (harvesting location) or biological (pathophysiologic reason for an increase/decrease in a cell type).

An asset of the GTEx project was the presence of digitized tissues for all tissues harvested (except specific brain regions), which was leveraged to validate cell-composition clusters. These clusters were essentially of two types – inadvertent cell types or structural differences in cellular composition. We were able to correlate 46 different clusters with histologic appearances of the tissue, using the extreme Z-score values of each cluster.

Two examples of variable clusters caused by inadvertent harvesting of adjacent tissues were exemplified by the coronary artery and the esophagus muscularis tissues. Cluster ARTCRN.F, in the coronary artery, represents adjacent accidentally harvested cardiac myocardial tissue collected during procurement and notable for 21 cardiac muscle genes including *MYH7*, *MYL3*, *TNNT2*, and *XIRP2*(**Fig. 3A**). Similarly, ESPMSL.B is a contaminating epithelial cluster (*KRT5*, *KRT4*, *MUC21*) in what should be exclusively esophagus muscular tissue (**Fig. 3B**).

**Figure 3.**
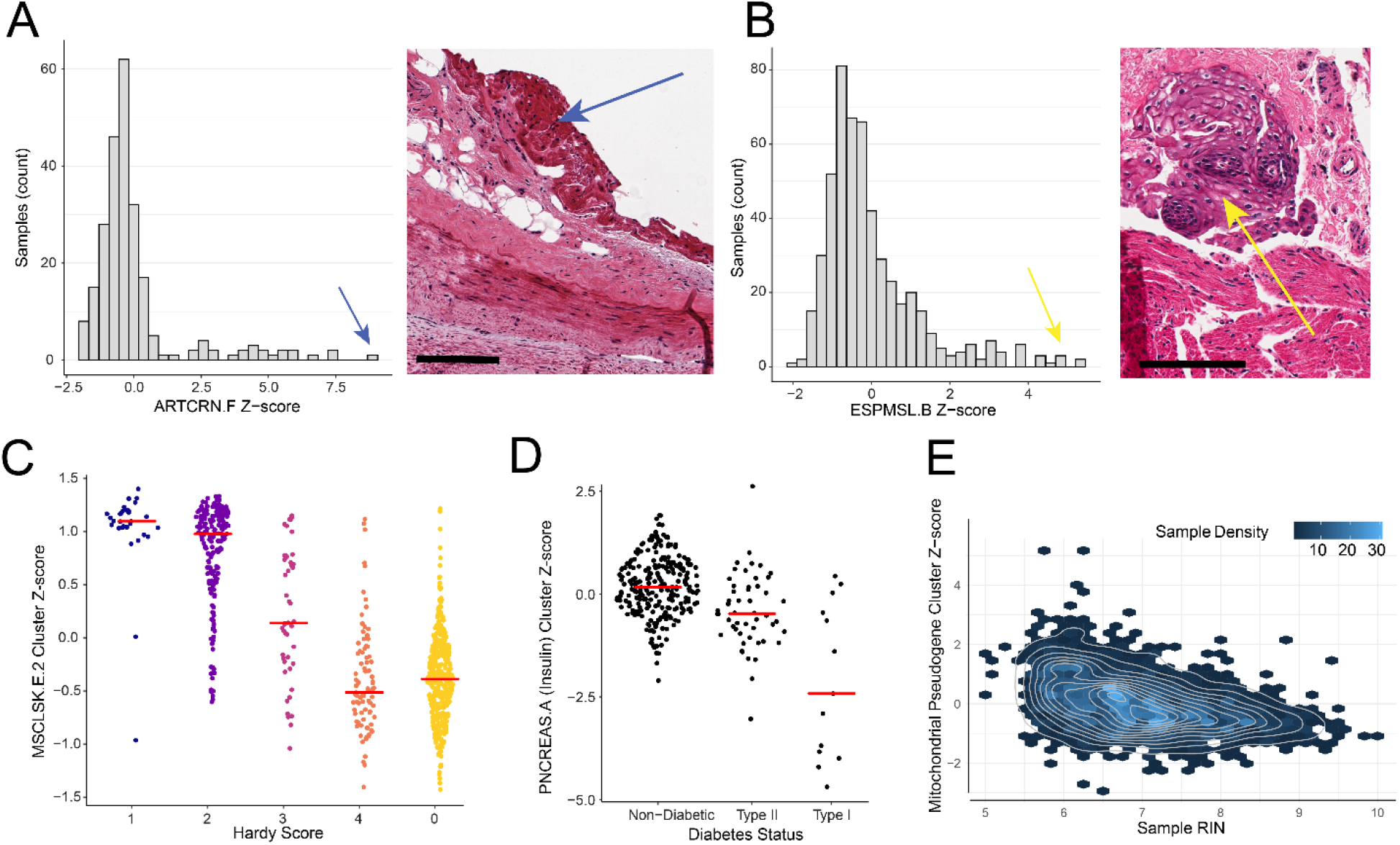
Unexpected and biologically driven organ specific clusters. **(A)** A histogram of the coronary artery, ARTCRN.F, cluster’s Z-score. This cluster is comprised of cardiac muscle markers (e.g. *MYH7*, *MYL3*, *TNNT2*) indicating heart contamination. A representative histology image of a coronary artery with cardiac tissue (blue arrow) corresponding to a sample with a high Z score. The bar is 150 μm. **(B)** A histogram of the esophagus muscularis, ESPMSL.B, cluster’s Z-score (blue arrow). This cluster includes epithelial cell markers (e.g. *KRT4*, *KRT19*). A representative histology image of esophagus muscularis with mucosa (yellow arrow) corresponding to a sample with a high Z score (yellow arrow). The bar is 150 μm. **(C)** A sina plot of the change in skeletal muscle, MSCLSK.E.2, Z-score across Hardy Score. Red bar = median value of the Z-score. **(D)** A sina plot representing the change in expression for cluster PNCREAS.A, which includes *INS*, across different diabetic types. Red bar = median value pf the Z-score. **(E)**A hexplot overlaid with a density plot comparing RNA Integrity Numbers (RIN) to the Z-scores of associated mitochondrial pseudogene clusters in all brain tissues that include a mitochondrial pseudogene cluster.

Sampling likely contributed to many other compositional clusters including the binary relationship between mucosa and submucosa (stroma/muscularis) in many gastrointestinal organs. An example of this is two negatively correlating clusters (CLNTRN.A and CLNTRN.C) of the transverse colon (**Supplemental Fig. S51**). CLNTRN.A (N = 199 transcripts), is an epithelial cell cluster, which includes the well-known epithelial cytokeratin marker gene *KRT20*(Chan et al. 2009). CLNTRN.C (N = 8 transcripts), a submucosa cluster, contains the smooth muscle marker gene *ACTA2* (Kendall’s Correlation; τ = 0.853; p-value = 2.39e-116). Additional composition-based clusters validated by histology include cluster BREAST.C, containing fat genes*LEP*and *ADIPOQ*(**Supplemental Fig. S52**). The most histologic adiposity among samples was seen with the highest Z-scores for that cluster. Similarly, MSCLSK.G, a skeletal muscle cluster containing genes *TNMD*and *COL22A1*, had high Z-scores among samples with tendinous tissue noted by histology (Docheva et al. 2005) (**Supplemental Fig. S52**.) A third cluster, HTRAA.D (heart, atria), notable for *KRT7*and *MSLN*, had increased mesothelial cell lining of tissues by histology among the high Z-score samples. (**Supplemental Fig. S52**). Some transcript clusters, which were likely caused by variable tissue elements, such as a cluster of hair-keratin associated genes (e.g.

*KRT75*, *KRT81, KRT82, KRT8*)*3*in skin (SKINS.B), could not be associated with the histology (Schweizer et al. 2006). This is likely due to the heterogeneity of some compositional elements. We had previously described this in lung and coronary artery tissue (McCall et al. 2016; Brehm et al. 2022) and note again how extreme Z-scores in lung cluster LUNG.A, **(Supplemental Table S2**), characterized by genes for cilia (e.g. *SPAG17*, *DNAH12*) and mucin (e.g. *MUC4*, *MUC5AC*) did not appear consistently by histology. These types of clusters indicate the importance of careful and consistent harvesting of tissues of interest.

### Disease related clusters: Skeletal muscle atrophy

A previous GTEx study indicated skeletal muscle was the tissue with the most genes affected by ischemic time (Ferreira et al. 2018). This was validated by our study, as 4 skeletal muscle transcript clusters, containing 149 total transcripts, associated with ischemic time. One of the clusters, skeletal muscle cluster E.2 (MSCLSK.E.2) containing 93 transcripts, positively correlated with both ischemic time and Hardy Score, but had a stronger correlation with Hardy Score (**Fig. 4C**).

**Figure 4.**
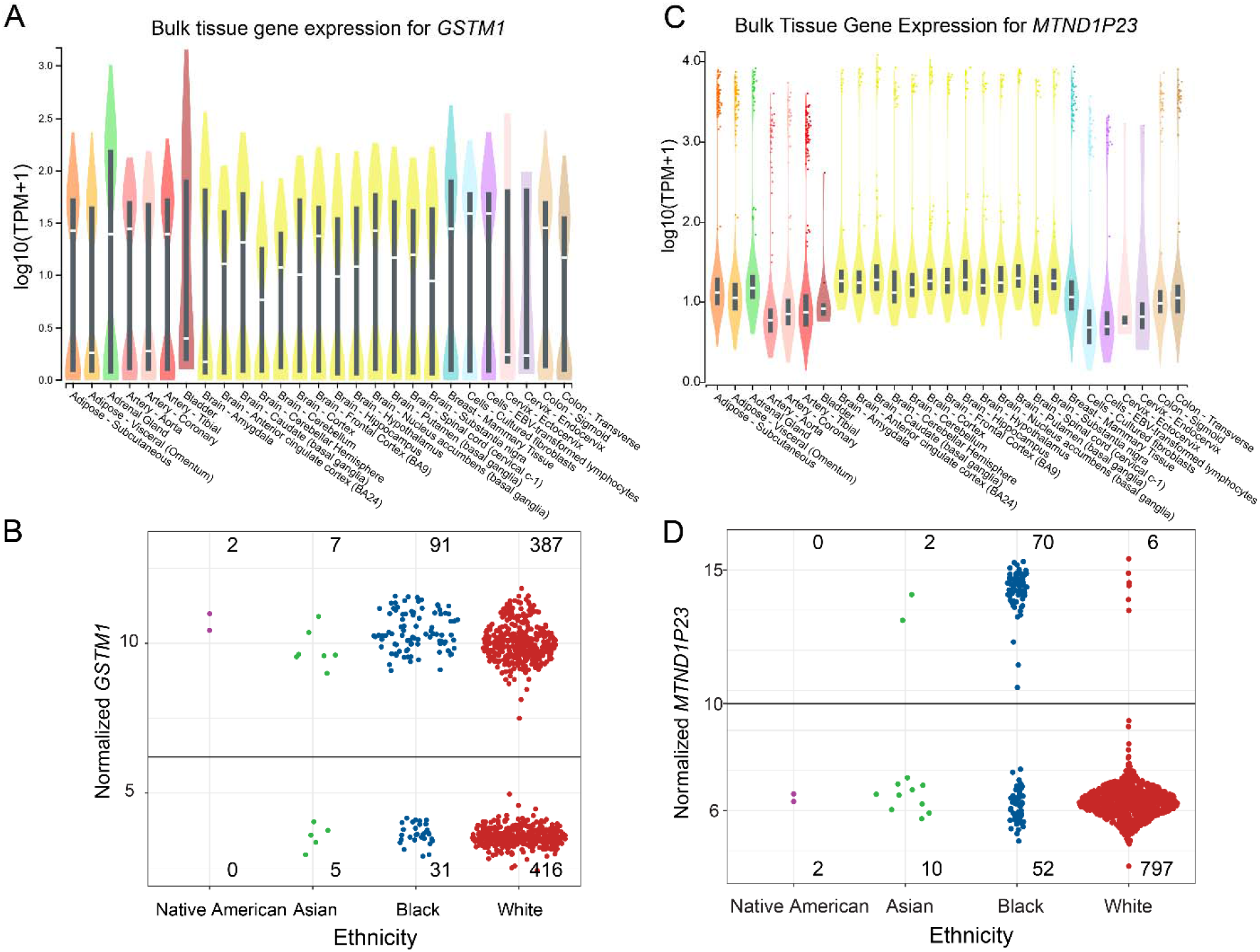
Extreme variability and population-based expression of transcripts. **(A)** Violin plot of *GSTM1*expression showing bimodal expression (from the GTEx Portal) **(B)** A sina plot of median normalized expression for *GSTM1*in 939 individuals by race. The Y-axis is variance stabilized transformed*GSTM1*values. **(C)** The pseudogene *MTND1P23*shows bimodal expression, but from fewer samples (from the GTEx Portal). **(D)** Median normalized expression for *MTND1P23*in 939 individuals by race, showing highest expression among black subjects. The Y-axis is variance stabilized transformed *MTND1P23*values.

We hypothesized that Hardy Score values, in the setting of prolonged hospital bed rest, could be a surrogate for inactivity and muscle atrophy. We therefore compared skeletal muscle clusters with genes known to be altered in critical illness myopathy (CIM) (Zink et al. 2009). A prior investigation of CIM identified 5,237 upregulated and 1,020 downregulated genes (Llano-Diez et al. 2019). We noted 48 of all 93 MSCLSK.E.2 transcripts (52%) or 48 of 78 genes (62%) were present among the CIM downregulated transcripts, which was highly significant (Fisher’s Exact Test; P-value = 4.99e-17). In the inversely correlated cluster MSCLSK.E.1, 8 of 38 transcripts (21%) or 8 of 28 genes (29%) were found in the upregulated CIM gene set. No E.2 transcripts were present among the 5,237 upregulated CIM genes and no E.1 transcripts were present in the 1,020 downregulated CIM genes. Interestingly, we also found 10 of 17 transcripts (59%) or 10 of 15 genes (67%) in the MSCLSK.J cluster were exclusively present in the upregulated genes (Fisher’s Exact Test; P-value = 2.04e-05). These clusters indicate the importance of identifying subject activity relative to tissue collection.

### Disease related clusters: altered islets in diabetes

In the pancreas, cluster A (PNCREAS.A), was notable for the presence of the insulin gene (*INS*). *INS* is made exclusively by beta cells of Islets of Langerhans. Glucagon *G*( *CG*), a hormone generated by Islet of Langerhans alpha cells was also variable, but not found in the PNCREAS.A cluster, even though it had a direct modest correlation with *INS,*(0.33 Kendall correlation, p-value: 5.87e-19). This suggested that a variable number of islets of Langerhans, per harvested sample, was unlikely to explain the PNCREAS.A cluster. Participant phenotype information indicated that 66 of 328 pancreas samples were obtained from type I or type II diabetic subjects. A linear model including diabetes status, age, sex, and tissue ischemic time, identified that diabetic status resulted in significantly lower Z-scores for the PNCREAS.A cluster, which was true in both type I and type II diabetic subjects ( **Fig. 3D**; linear model; type I beta = -2.31, p-value : 1.16e-21; type II beta = -0.60, p-value: 5.92e-06). Without diabetic subjects, the PNCREAS.A transcript group would not be sufficiently variable to be listed as a cluster (variance with diabetic subjects = 3.65; variance without diabetic subjects = 0.81), indicating the importance of identifying participant disease status.

### Harvesting related clusters: Brain, mitochondrial pseudogenes and pH

A cluster of variably expressed mitochondria pseudogenes and tRNA transcripts (including *MT-TY*, *MT-TE*, *MT-TP*, *MTND1P23*, *MTND2P28*and *MTCO1P12*) were present in 11 separate brain region clusters and appeared as unclustered variable transcripts in an additional 2 brain regions. We hypothesized this was a technical batch effect. An initial linear mixed model containing 37 variables and 2 random effects (tissue and subject ID) strongly identified the intron-related variable ‘split reads’ (SMSPLTRD: split reads; beta: -0.61; p-value: 1.15e-35) (**Supplemental Table S4**). All intron-related variables were removed from the model as mitochondrial pseudogenes/tRNAs do not have introns. In a filtered model, the most variance was explained by the RNA integrity number (SMRIN, linear mixed model; beta: -0.11; p-value: 1.13e-10; **Fig. 3E; Supplemental Table S**).**5**Additionally, the variables ‘number of rRNA alignments,’ ‘brain pH,’ and ‘alternative alignments’ were also significant (p-values: 6.07e-9, 3.04e-6, and 6.94e-5 respectively). Overall, this analysis indicated that the mitochondrial pseudogene/tRNA transcript clusters are a feature of tissue procurement and quality, but not biological variation. The cluster indicated the importance of capturing RNA quality metrics on collected tissues.

### Harvesting related clusters: Premortem blood draw vs postmortem blood draw

Blood cluster B, had two sets of inversely correlated transcripts (BLOOD.B.1 with 63 transcripts, BLOOD.B.2 with 104 transcripts). Each cluster had a near bi-modal Z-score distribution based on the blood being acquired pre or postmortem ( **Supplemental Fig. S53**, Wilcoxon Rank Sum test B.1 p-value = 1.81e-94; B.2 p-value = 3.18e-111). Blood procurement was a feature of a patient being hospitalized (premortem) or dying acutely (postmortem). Previous research using the same GTEx dataset has shown that large drivers of variation between pre and post-mortem blood draws are hypoxia and immune cell type (Ferreira et al. 2018).

### Age related clusters: Menopause and the uterus

We had hypothesized that the uterus would undergo aging/menopausal changes that could result in transcription variation. We investigated this relationship among the 22 uterus clusters, many of which were modestly correlated to each other. Menopause was not recorded by GTEx, so we binned the donors into three separate groups based on the age quartiles (premenopausal first quartile, perimenopausal second two quartiles, postmenopausal fourth quartile) (McKinlay 1996). We observed that four clusters, UTERUS.G, UTERUS.J.1, UTERUS.A.1 and UTERUS.M.1 collectively had more lowly expressed Z-scores with higher age quartile (**Supplemental Fig. S54,** linear model p-value = 1.49e-14; beta = -0.89). This indicated an age/menopausal related reduction of expression for these gene sets.

### Nonclustering, strongly variable transcripts

Despite the strong clustering of most transcripts, several transcripts were commonly variable across tissues, but did not cluster/correlate in expression with 5 or more other transcripts. Of the 4,961 variable transcripts, 2,711 were present in two or more tissues. Overall, the most variable transcripts were *GSTM1*, *RP11-343H5.4*, *NPIPB15*, *MTND1P23* and *C21orf33* which, despite being variable in 43-49 tissues ( **Supplemental Table S2**), did not cluster with other transcripts (outside of the brain mitochondrial pseudogene cluster). *GSTM1* and *NPIPB15* are known to have population-variable null alleles (McIlwain et al. 2006) while *RP11-343H5.4*is a pseudogene transcript known to have genotype dependent alignment variability at different frequencies across human populations (dbSNP rs4841) (Saha and Battle 2018) ( **Fig. 4A-B**.) *MTND1P23* has a SNP common in Africans (∼50%), rs6594028, that allows perfect alignment for *MT-ND1* reads (Fisher’s Test p-value: 8.06e-64 and 5.95e-09 respectively; **Fig. 4C-D**.) *C21orf33* appears to have null alleles in Caucasian samples. Collectively, these transcripts showed extreme, bimodal expression across samples, rather than more traditional distributions, which may cause them to be identified as widely variable for non-disease related reasons in genomics-based studies.

### Applicability to single cell sequencing studies

All of these biological and technical drivers of variation were described in bulk sequencing. We questioned the applicability of these findings to single cell sequencing data, where depth of sequencing would make this type of study challenging, and applied these discoveries to Tabula Sapiens data available through CellxGene (Megill et al. 2021; Tabula Sapiens et al. 2022). We noted many of our discoveries were important potential drivers of clustering variability and discoverable as major variable genes.

One driver of variation that we describe here, and have reported on previously, is sequencing contamination. We already noted its impact on the Tabula Muris data (Tabula Muris et al. 2018; Nieuwenhuis et al. 2020). We again note that cross sample contamination, well-described in GTEx also causes cell-cell cross contamination in single cell data. For example, liver endothelial cells clustered away from the main endothelial cell cluster ( **Fig. 5A-C**). Many of the genes highly expressed (in a relative fashion) in these endothelial cells, are very highly expressed genes in hepatocytes ( *SAA1*, *SAA2*, *HP*) and that ambient RNA cross-contamination is adding these transcripts to the endothelial cell counts, causing them to cluster away from other endothelial cells.

**Figure 5.**
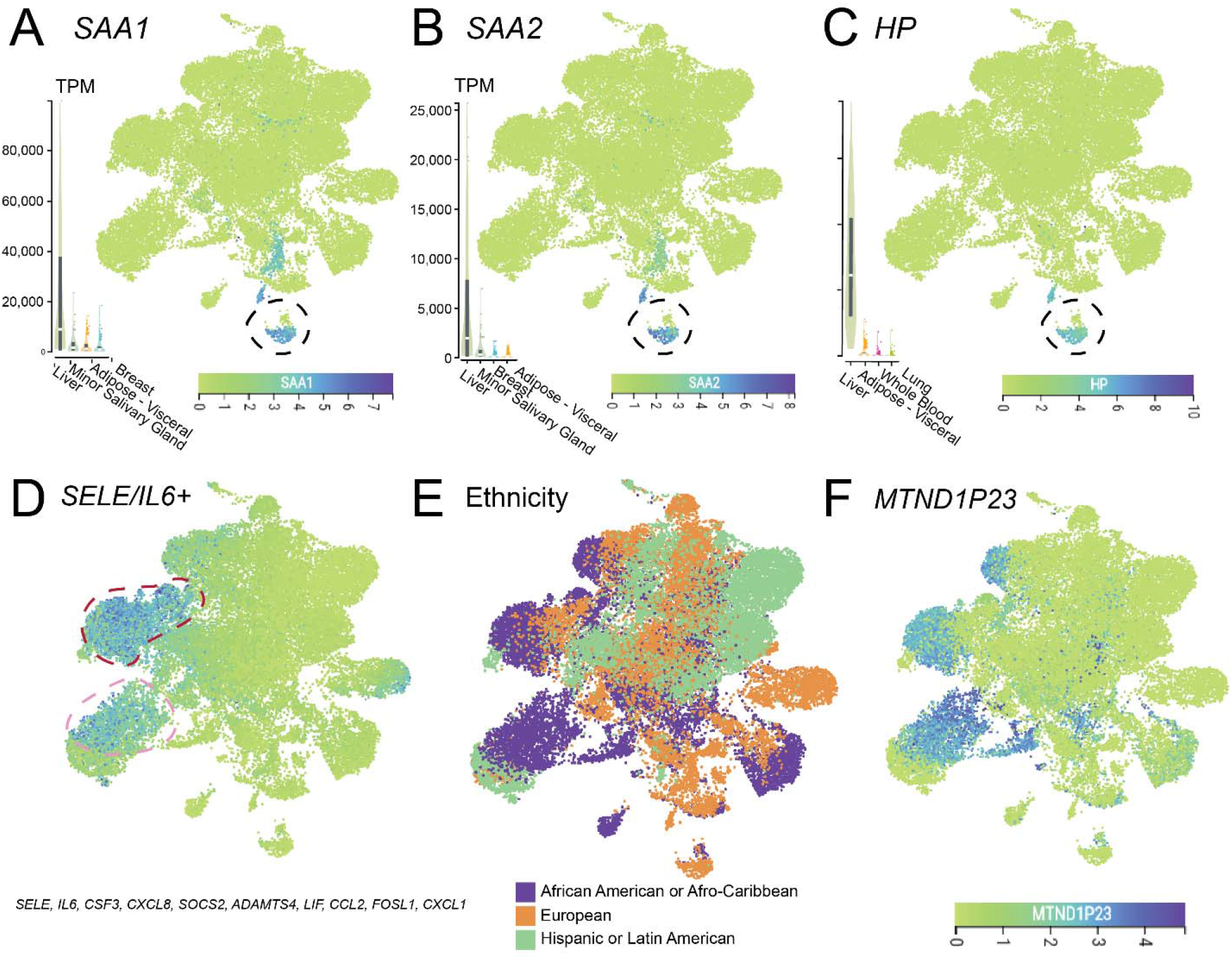
Impacts of variation diversity on single data in clustered Tabula Sapiens endothelial cells. **(A-C)** A UMAP of endothelial cells where liver endothelial cells (circled in black) are contaminated by highly expressed hepatocyte-specific genes. Dark blue indicates high expression. Transcripts per million (TPM) values, sorted from highest to lowest are shown for the 4 top tissues from the GTEx portal. **(D)** High expression of the endothelial activation cluster in specific subsets of endothelial cells. **(E)** The distribution of ethnicities in the Tabula Sapiens endothelial population, showing the activation cluster was seen in only a subset of samples. **(F)**Expression of *MTND1P23*, a pseudogene with expression enriched in groups of African heritage, is noted specifically in samples from African American or Afro-Caribbean descent.

Another driver of variability we described was of endothelial activation ( *IL6*/ *SELE* cluster) associated with the speed of the final stage of death (Hardy Score). A search of this gene set (from **Fig. 2E**) in the endothelial cell Tabula Sapiens UMAP identified two localized positive clusters of cells (**Fig. 5D**). This cluster was elevated in one muscle sample and one uterus sample, suggesting these donors likely had different death characteristics than other samples.

Finally, we note that a strength of Tabula Sapiens is that the samples are derived from multiple donors across different ethnicities. We showed in GTEx data that expression of the mitochondrial pseudogene *MTND1P23,*is the result of misaligned *MT-ND1*reads in Black individuals with a rs6594028 SNP G allele. This pseudogene highlights specific cell clusters from individuals labeled as African-American or Afro-Caribbean in CellxGene (**Fig. 5E-F**). While its presence/absence did not alter cell clustering, a comparison of “endothelial cell of vascular tree” cells which came from African-American/Afro-Caribbean or European subjects identified *MTND1P23* as the most variable gene across the samples. Thus, drivers of variation uncovered in the bulk sequencing GTEx dataset are likely applicable to single cell studies.

## DISCUSSION

This is the first large scale survey of drivers of expression variation in human tissues. We uncovered 522 clusters of variable genes across 49 tissues. Unsurprisingly, both biological and technical factors caused variation. Among the technical factors, the common causes included sequencing contamination, ischemic time, and brain pH. Among the biological factors, the most common causes were death interval and tissue variability.

Another large driver of variation was tissue composition variability due to differences in sampling. This sampling challenge could be the result of biological or technical factors. Among biological reasons are subtle organ structural differences in size and shape between subjects such as different thicknesses of renal cortex or adrenal medulla. These can be inherited, a result of normal biological processes, or could reflect an acute or chronic response to disease. Among technical factors are differences between procurement teams where one group may obtain a tissue section more medial or lateral to another team, may or may not trim excess tissues such as adipose off specimens as completely, or may wash away blood differently. Nearly every tissue had evidence of cell type composition as a cause of at least one variable gene cluster.

The largest surprise to us, of drivers of variability, were the two inversely related clusters associated with Hardy Score. Although cause of death is frequently obtained from death certificates, the time course of disease and ventilator use are not. Thus, this terminal death phase, captured by Hardy Score, would be a useful phenotypic datapoint to be collected going forward for tissue-based research studies.

Many tissues had unique causes of variation. We highlighted several examples, but many more were noted (Supplemental Table clusters). These often related to variable amounts of inflammation, the capture of different brain regions/nuclei, organ-specific diseases, or responses to therapeutics.

It is almost certain that many clusters of variable genes exist together due to multifactorial causes and that our identified causes may represent only some of the drivers of variation. As an example, the cluster of lung genes that included surfactant proteins and indicated ventilator associated lung injury, is also associated with a shorter postmortem interval, as ventilated patients had the shortest time to autopsy. This is an example of how many metadata elements correlate with each other. Additionally, many (n= 74) clusters could not be explained by any of the available metadata data. While no study can capture all phenotypic elements, some datapoints that were missing may have been relevant for particular variation clusters. As an example, knowledge of the medications of each subject may have helped solve liver cluster LIVER.A.1, containing cytochrome P450 genes, which can become upregulated depending on medication usage (Luo et al. 2004). Thus, even the GTEx dataset, with some of the most extensive metadata of any large study still has gaps in the information it captured and recorded.

There were a number of limitations of the project. Several clusters that suggested cellular composition differences could not be validated in the available histologic images. This could be due to heterogeneity of the sampling of the piece used for histology vs. the piece used for RNA studies, as we described (Brehm et al. 2022). Also, there was no matching histology for the brain sections, which precluded this analysis for 13 tissues. It is unclear the generalizability of these variation clusters to other studies. To determine this, one will need to investigate other, more tissue-limited projects and note if these same gene clusters appear. Certainly, the gene composition of contamination clusters will depend on what other tissues are being prepared and sequenced concurrently. Of note, the presence of the endothelial activation cluster ( *SELE*, *IL6*, etc.) in single cell sequencing projects (**Fig. 5d**) does suggest some of these clusters are likely to be universal phenomena (Reichart et al. 2022). Finally, we used high gene variance as a convenient method to detect genes that differed across samples of the same tissue; however, we have missed some genes with a smaller magnitude of expression changes between the tissues that still represent an important biological or technical source of variation. Future studies could seek to relax the variance threshold used in this work or implement alternative approaches to identify genes whose expression differs between samples from the same tissue.

In conclusion, both biologic and technical factors contribute to tissue expression diversity. Using the large and robust GTEx dataset, we were able, for the first time, to describe and identify numerous causes of tissue-based expression diversity. We expect this data can inform on better approaches to normalize datasets where a more active approach, rather than latent factors, can be used, particularly when specific variable phenotypes (e.g. disease) are important to the analysis.

## METHODS

### Acquisition of GTEx Transcriptomic and Phenotype Data

The raw read count file for all GTEx samples (GTEx_Analysis_2017-06-05_v8_RNASeQCv1.1.9_gene_reads.gct.gz) was acquired from the GTEx portal website (https://gtexportal.org/home/datasets; last accessed: 3/9/2021). Deidentified sample data was also acquired through the portal (GTEx_Analysis_v8_Annotations_SampleAttributesDS.txt), while individual phenotype data was acquired through dbGap with approval. Pertaining to human data, all methods were carried out in accordance with relevant guidelines and regulations.

### Processing of bulk sequencing data and establishment of variable genes

Each tissue was separately processed through the following pipeline in R (4.1.1). All gene expression levels were normalized using DESeq2’s VarianceStabilizingTransformation function (1.34.0). Genes which had an across sample mean value of < 5 normalized counts were removed. The variance of each remaining gene was calculated and those in the top 2% for each tissue became our high variance gene dataset.

### Correlations and hierarchical clustering using both agglomerative and divisive approaches

Correlations were calculated for the high variance gene sets using the Kendall’s τ method. One minus the absolute value of this correlation matrix was used as the distance in a hierarchical dendrogram created using the base R function hclust() with the “average” linkage method. To produce subgroupings based on the hierarchical clustering, we generated a Kendall’s τ critical value from a function of the inverse of a Kendall’s τ distribution, the probability being calculated as below, where the number of genes is denoted as *ngene,* while the number of treatments for the distribution function being the number of samples.

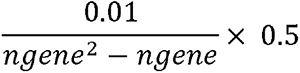

Using base R’s cutree() function the generated dendrogram was cut at one minus the Kendall’s τ critical value giving an initial set of clusters. These clusters were then filtered to the subset composed of 6 or more genes.

An agglomerative approach then evaluated the need to subdivide clusters into potentially more specific components. A global threshold correlation value for all tissue high variance genes was set at the 80th percentile of all high variance correlations. If the mean of the absolute values of a given cluster were below the 80^th^ percentile cutoff, that cluster was further evaluated for subdivision.

For the given cluster, a 75^th^ percentile threshold value is set based on the correlations of the cluster genes. Then the cluster is broken into individual genes and gene clustering is performed in a stepwise fashion from a 100% correlation (correlation of a gene only to itself) down to the 75 ^th^ percentile threshold value. Each separate subcluster becomes a unique gene cluster for further analysis and the original larger cluster is removed. These new clusters also have a ≥6 gene requirement to be evaluated. Finally, recognizing a ‘one-size-fits-all’ approach for clustering all tissues may result in some inappropriate cluster profiles, a manual review was performed which modified ∼20 clusters where too fine of separation yielded excess clusters. All clusters are named by the GTEx tissue abbreviations and sequential alphabetical letters (ex. SKINS.B) Lastly, clusters are subdivided if they have genes that negatively correlate together within them, eg. *XIST* and Y-chromosome genes (ADPSBQ.G.1, ADPSBQ.G.2).

### Cluster normalized Z-scores

Z-scores were created for each cluster to summarize their relative expression in separate GTEx samples. The equation is below, where *x* is the VST normalized expression of gene *j* in sample *i*, *t(i)* is the tissue type for sample *i*, and *J* is the number of genes in a given cluster .

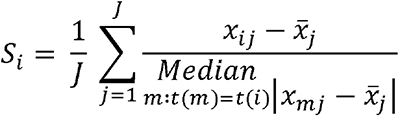

### Hemoglobin analysis

We extracted VST normalized values for*HBB*, *HBA1, HBA2,* and *HBD*in the following tissues ADRNLG, ARTAORT, CLNSGM, CLNTRN, ESPGEJ, ESPMCS, ESPMSL, HRTAA, HRTLV, SLVGLND, STMACH, and TESTIS. These tissues were used in a previous ischemic time analysis (Ferreira et al. 2018). For each hemoglobin gene we ran a separate linear mixed model (lme4 v1.1-27.1) including ischemic time, age, height (HGHT), weight (WGHT), body mass index (BMI), ethnicity (ETHNCTY), RNA integrity number (SMRIN), sample center (SMCENTER), sample batch (SMNABTCHD), cohort (COHORT), tissue (SMTSD), with subject ID (SUBJID) as a random effect. Covariates were selected based on previous similar studies (Ferreira et al. 2018).

To generate a hemoglobin score, we used the cluster normalization method on the previous listed tissues on *HBB*, *HBA1, HBA2,*and *HBD*. We then reran the above model with the outcome being the new hemoglobin score.

### Key metadata variables

Hardy Score (DTHHRDY) is defined as the terminal death interval and is measured as an ordinal variable ranging from 0 to 4 with the following definitions: 1 = Violent fast deaths with a terminal phase estimated at < 10 min. 2 = Fast death of natural causes with a terminal phase estimated at < 1 hr. 3 = Intermediate death after a terminal phase of 1 to 24 hours 4 = Slow death after a long illness, with a terminal phase > 1 day. 0 = All cases on a ventilator immediately before death.

GTEx has two separate ischemic variables: individual ischemic times (TRISCHD: GTEx Procedure Start Time Interval between actual death, presumed death, or cross clamp application and the start of the GTEx Procedure in minutes), and sample ischemic times (SMSTISCHD: Total Ischemic time for a sample in minutes). Unless stated otherwise in the text, all ischemic times used in this analysis are sample (SMSTISCHD).

Digital slides of tissues were acquired directly through the GTEx portal’s histology viewer (https://gtexportal.org/home/histologyPage; or https://brd.nci.nih.gov/brd/image-search/searchhome last visited 12/28/2021). Other metadata variables used in this study are only available through DBGap permissions. These include, but are not limited to, unbinned age (AGE), race (RACE), diabetes status (MHT1D and MHT2D), death on ventilator (DTHVNT), and brain pH (MHBRNPH).

### Tools used to solve cluster identities

In evaluating individual clusters of genes, we utilized a number of online resources that could indicate shared functionality or cell-type localization. These included CellxGene [https://www.biorxiv.org/content/10.1101/2021.04.05.438318v1], StringDB (Szklarczyk et al. 2021), Gene Ontology (Ashburner et al. 2000; Gene Ontology 2021) pantomir, and HPAStainR (Nieuwenhuis and Halushka 2020). Gene lists were entered into the various software tools either manually (CellxGene, StringDB) or via pipelines (HPAStainR). Significant findings were cross-validated by GeneCards, OMIM, or a literature search of the key genes or pathways identified. A general understanding of each cluster’s gene composition was then cross-referenced with hypothesized reasons for variability of the cluster. A number of testable hypotheses were performed with the given metadata resources from GTEx. This included comparisons to provided subject and sample technical factors (ischemic time, RIN, sample batch), biological factors (age, sex, Hardy score), genetic factors (SNPs, RACE), and digital images of tissues with the most extreme Z-scores by cluster.

### Computational methods to correlate clusters to metadata

#### Hardy Score cluster correlations

We used base R’s linear model function to build 81 linear models, one for each cluster including *PLA2G2A, CHI3L1, FOSB, SE*a*L*n*E*d/or *IL6*. The outcome for each model was the cluster’s Z-score with the predictors being sample ischemic time and Hardy Score coded as a linear ordinal factor with ventilator individuals being treated as 5s. Covariates in the model included age, sex, and sequencing batch. The analysis was only limited to the ischemic time and Hardy Score covariates which had their p-values corrected for using a Benjamini-Hochberg correction, significance noted as an FDR < 0.05.

### Skeletal Muscle Analysis

Using the sample data and individual phenotype data from the GTEx portal and the normalized Z scores for cluster MSCLSK.E.2 we built a linear model, where*h(i)* is the Hardy Score of the sample, *s(i)* is the sex of the sample, *a(i)* is the age of the individual the sample comes from, and *smisch(i)* is the ischemic time of the sample. Sex, age, and ischemic time were all included as they are the well-known batches (Somekh et al. 2019).

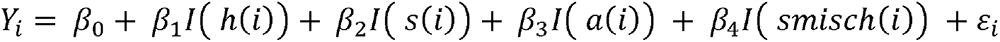

Up and downregulated genes in critical illness myopathy were acquired from additional file 2 of Llano-Diez et al. (https://static-content.springer.com/esm/art%3A10.1186%2Fs13395-019-0194-1/MediaObjects/13395_2019_194_MOESM2_ESM.txt; last accessed: 10/28/2021). A Fisher’s Exact Test was used to determine enrichment in high variance genes and MSCLSK.E.2.

### Pancreas Analysis

Phenotype data on type I and type II diabetes (MHT1D and MHT2D) was used to create a “diabetes” factor variable indicating if an individual was a type I diabetic, a type II diabetic, or a non-diabetic. We then created a model on the normalized PNCREAS.A values including diabetes, age, sex, and tissue ischemic time as covariates.

We performed the cluster analysis described above with and without diabetic individuals to evaluate *INS*variance. Starting at the raw counts, diabetic individuals based on the “diabetes” column were removed from the analysis. We then repeated the tissue analysis until the variance filter, where we saw *INS* could not pass the filter.

### Uterus Analysis

To create pre-, peri-, and post-menopausal bins the age of individuals with uterus samples were binned into quartiles. Premenopausal first quartile (age ≤ 39.25; N= 36), perimenopausal second two quartiles (39.25 > age < 58; N = 69) and a postmenopausal fourth quartile (age ≥ 58; N = 37) were established. A combined cluster Z-score was created for clusters UTERUS.G, UTERUS.J.1, UTERUS.A.1, and UTERUS.M.1 by taking the average Z-score value across these clusters for a given individual. The linear model was run with the outcome being combined cluster Z-score and the predictor being menopausal bin, with sample ischemic time as a covariate.

### Brain RIN Score Analysis

To investigate the 11 brain tissue mitochondrial pseudogene/transcription factor clusters, we used the normalized Z scores for clusters that included the following genes: *MTCO3P12*, *MTCO1P12*, *MTND1P23*, *MTND2P28*, *MT−TS1*, *MTND4P12*, *MT−TE*, *MTATP8P1*, *MTCO2P12*, *MT−TY*, *MT−TL2*, and *MT−TP*. A linear mixed model was performed which included the covariates found in **Supplemental Table S5**with both tissue and the SUBJID being random effects. We chose this model as it outperformed simpler models with only SMRIN, SMTSISCH, sex, age, and tissue with SUBJID as random effect via ANOVA and Akaike information criterion.

### Polymorphism Analysis

sVST normalized counts for*GSTM1*, *RP11-343H5.4*, *NPIPB15*, *MTND1P23*and *C21orf33* for 35 tissues and in total 12,917 samples were subsetted out. Then for each individual, we took the median expression of the transcript resulting in one value per person (N = 939 individuals). Individuals were binned on race (RACE) and an arbitrary threshold was created that best split the bimodal distribution. A Fisher’s Exact Test was used to determine if the distribution of expression was unexpected in contrast to race.

## DATA ACCESS

All datasets used in this manuscript are from previously reported studies available through the GTEx portal (https://gtexportal.org/home/) or dbGAP (https://www.ncbi.nlm.nih.gov/gap/) or through CellxGene (https://cellxgene.cziscience.com/collections/e5f58829-1a66-40b5-a624-9046778e74f5).

## COMPETING INTEREST STATEMENT

The authors declare that they have no competing interests.

## ACKNOWLEDGEMENTS

This work was supported by the National Heart, Lung, and Blood Institute (R01HL137811) to M.K.H., National Institute of General Medical Sciences (R01GM130564) to M.K.H., National Institute of General Medical Sciences (R01GM139928) to M.N.M., and the University of Rochester (CTSA, UL1TR002001) to M.N.M.

## SUPPLEMENTAL MATERIALS

**Supplemental Figs. S1 – S49:** Heat maps of 522 gene clusters across 49 GTEx tissues, based on Kendall’s τ. Correlations range from 1 (red) to -1 (blue) for genes.

**Supplemental Table S1.**Full names and abbreviations of GTEx samples used in the text and figures.

**Supplemental Table S2**. 522 cluster groups of genes across 49 GTEx tissues with cluster etiology and certainty.

**Supplemental Table S3**. Unclustered, variable genes across 49 GTEx tissues.

**Supplemental Table S4.**Initial linear mixed model containing 37 variables and 2 random effects (tissue and subject ID) for evaluation of brain pseudogene clusters.

**Supplemental Table S5.**Filtered linear mixed model containing 37 variables and 2 random effects (tissue and subject ID) for evaluation of brain pseudogene clusters.

---

**Supplemental Fig. S50.**
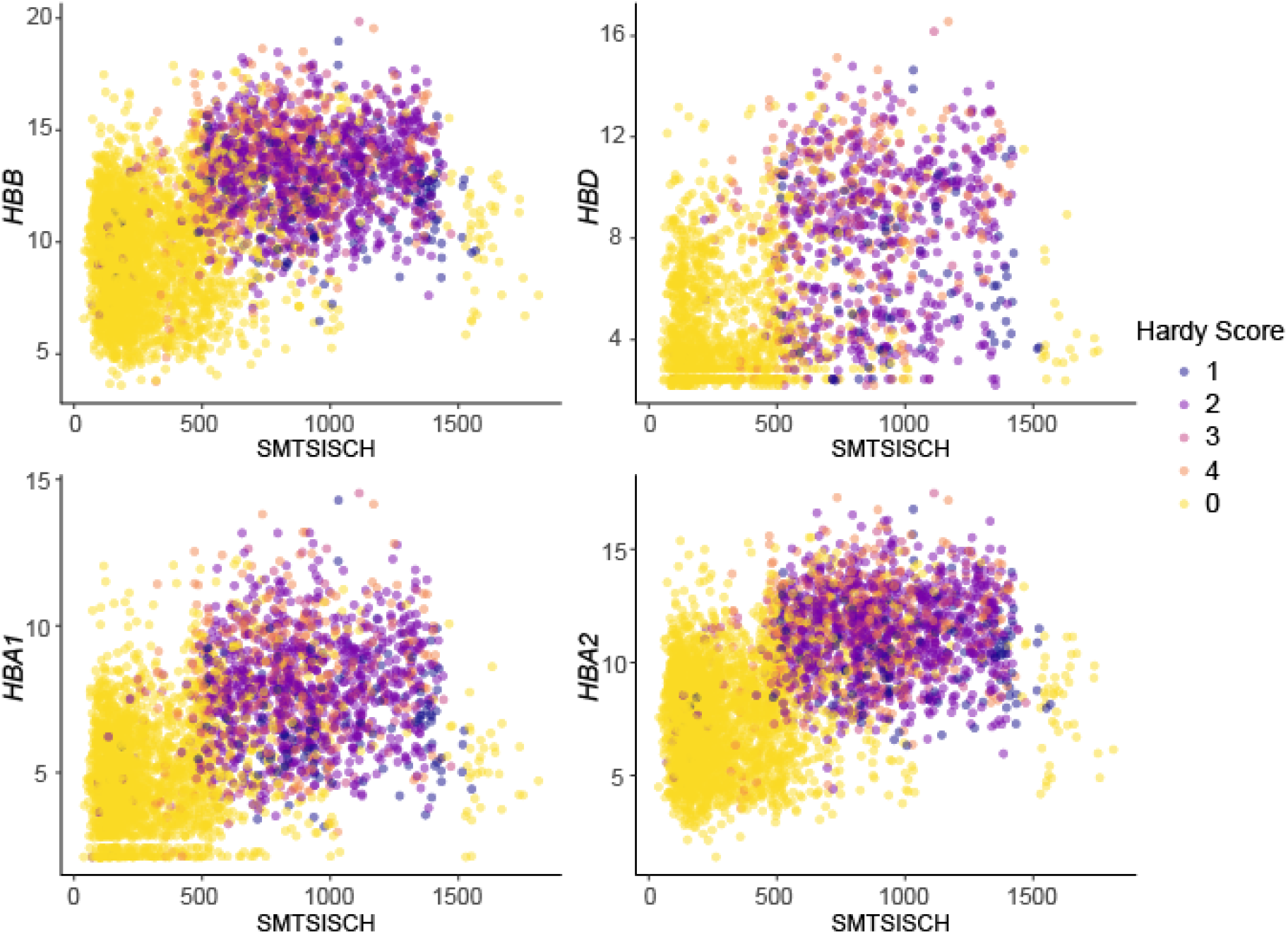
Relationship of hemoglobin genes to Hardy Factor. The longer the ischemic time (SMTSISCH), the higher the hemoglobin values. Longer ischemic time intervals associated with more acute deaths (Hardy score 1 and 2). Most short ischemic time intervals were in ventilated patients (Hardy score 0).

**Supplemental Fig. S51.**
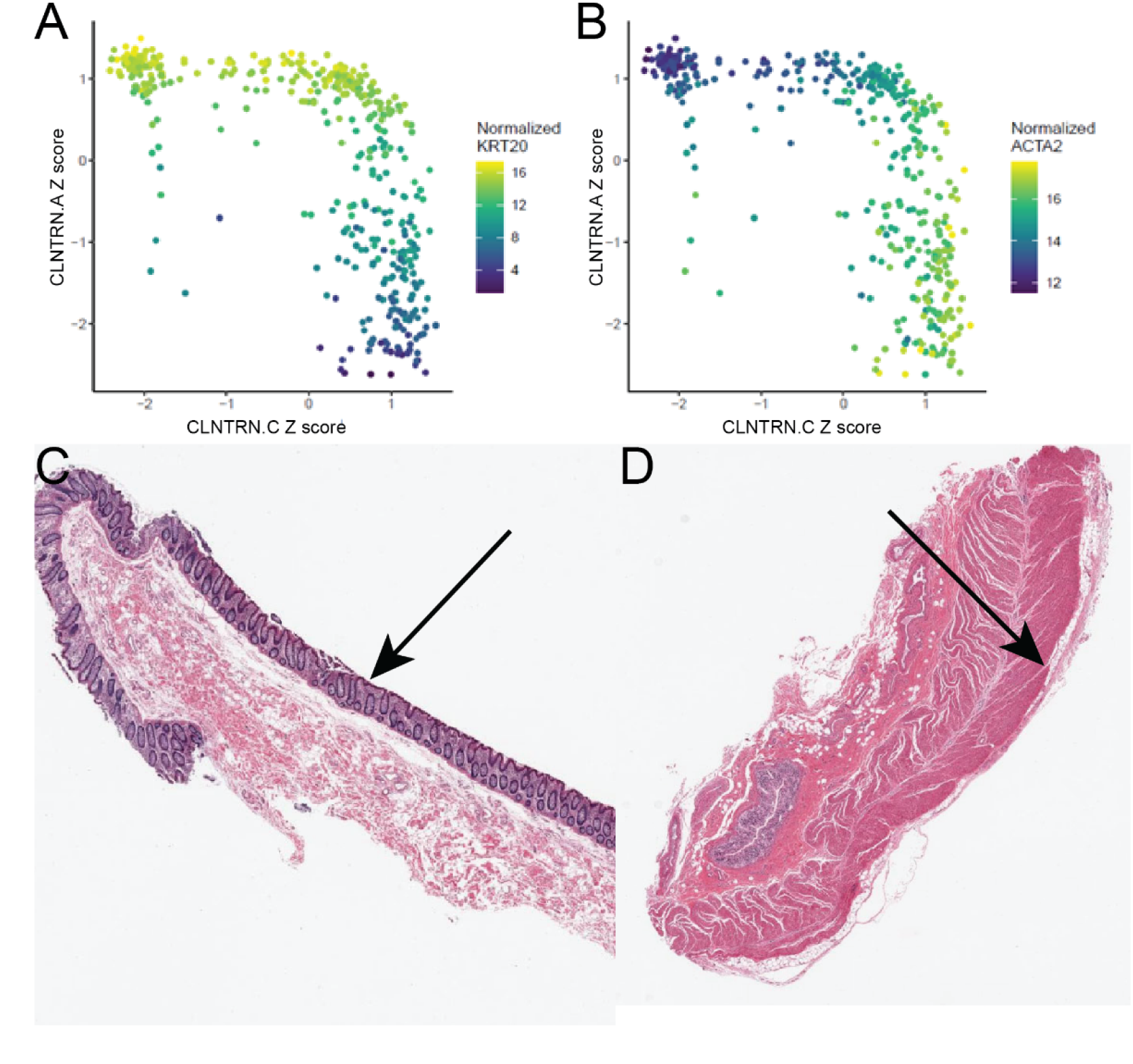
Transverse Colon illustrating the mucosa stroma dichotomy in the gastro-intestinal tract. **(A)** A scatter plot of CLNTRN.A’s Z-score on the Y-axis (mucosal genes) with CLNTRN.C’s Z-score on the X-axis (stromal genes) colored b*K*y*RT20* expression. **(B)** The same plot colored by *ACTA2*expression. **(C)** Histologic image of high CLNTRN.A Z-score demonstrating abundant mucosa (arrow) and an absence of underlying smooth muscle cells. **(D)** Histologic image of high CLNTRN.C demonstrating abundant smooth muscle cells (arrow) and absent mucosa.

**Supplemental Fig. S52.**
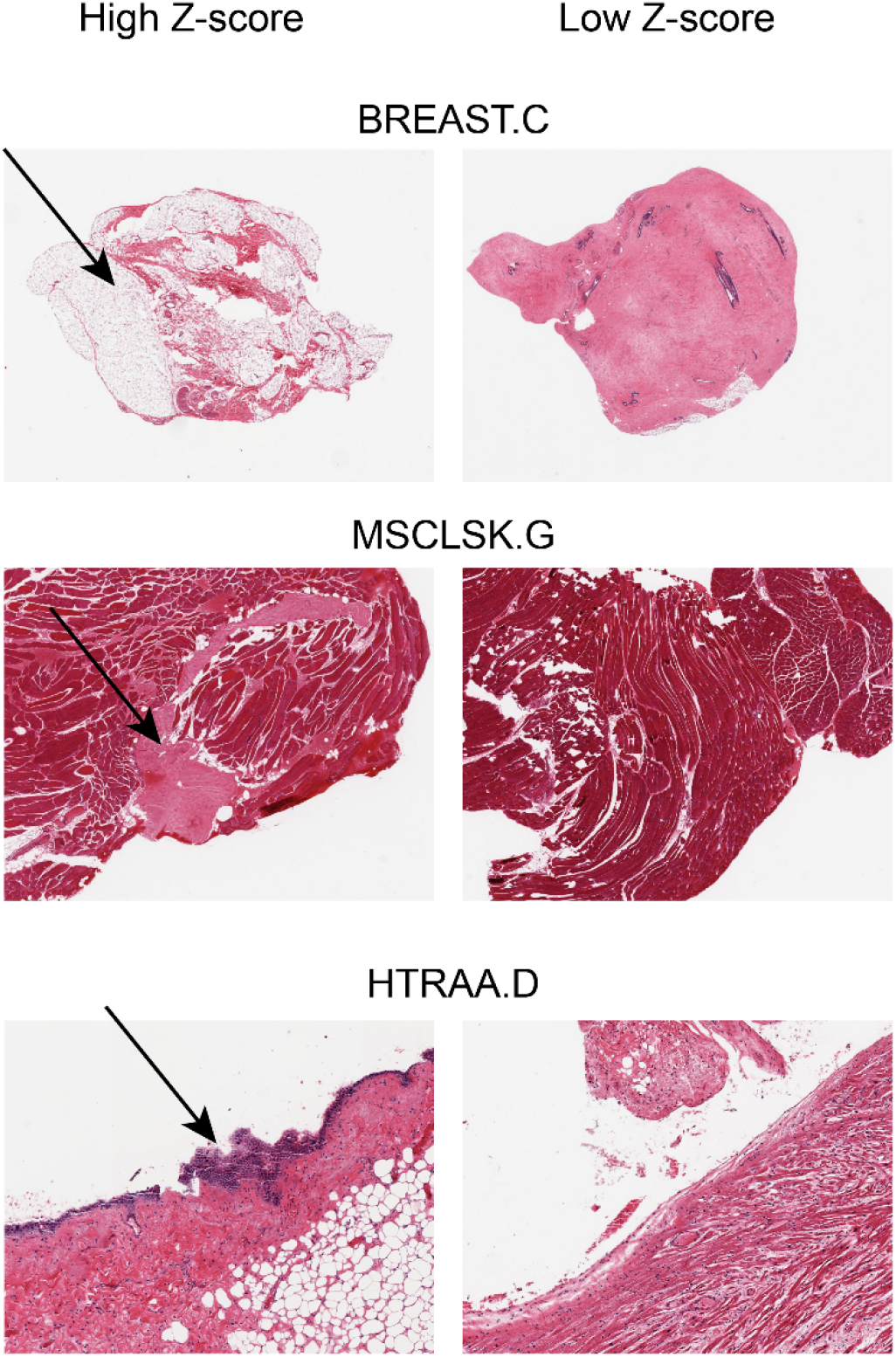
Histologic examples of cellular composition of tissues causing variation clusters. The BREAST.C cluster was associated with fat genes and shows extreme differences in fat content (arrow) between high Z-score samples (left) and low Z-score samples (right). The musculoskeletal cluster MSCLSK.G had high Z-scores correlating to the presence of tendon tissue (arrow, left) or absence of tendon tissue (right). The heart atrium (HTRAA.D) cluster indicated the presence (arrow) or absence of surface mesothelial cells.

**Supplemental Fig. S53.**
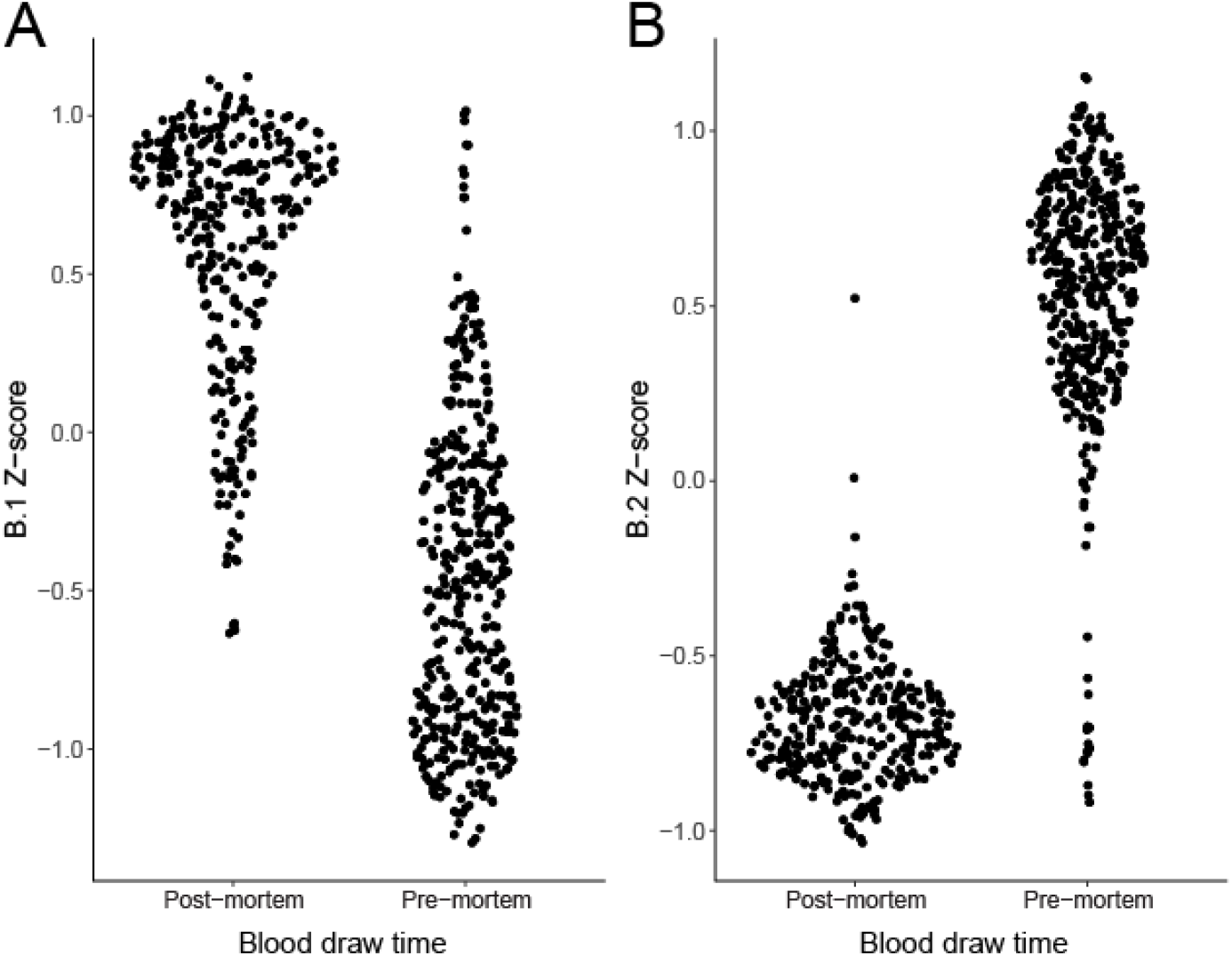
Pre and post-mortem blood draw clusters. **(A)** A sina plot of the BLOOD.B.1 Z-score binned on pre or post-mortem blood draw. **(B)** A sina plot of the BLOOD.B.2 Z-score binned on pre or post-mortem blood draw.

**Supplemental Fig. S54.**
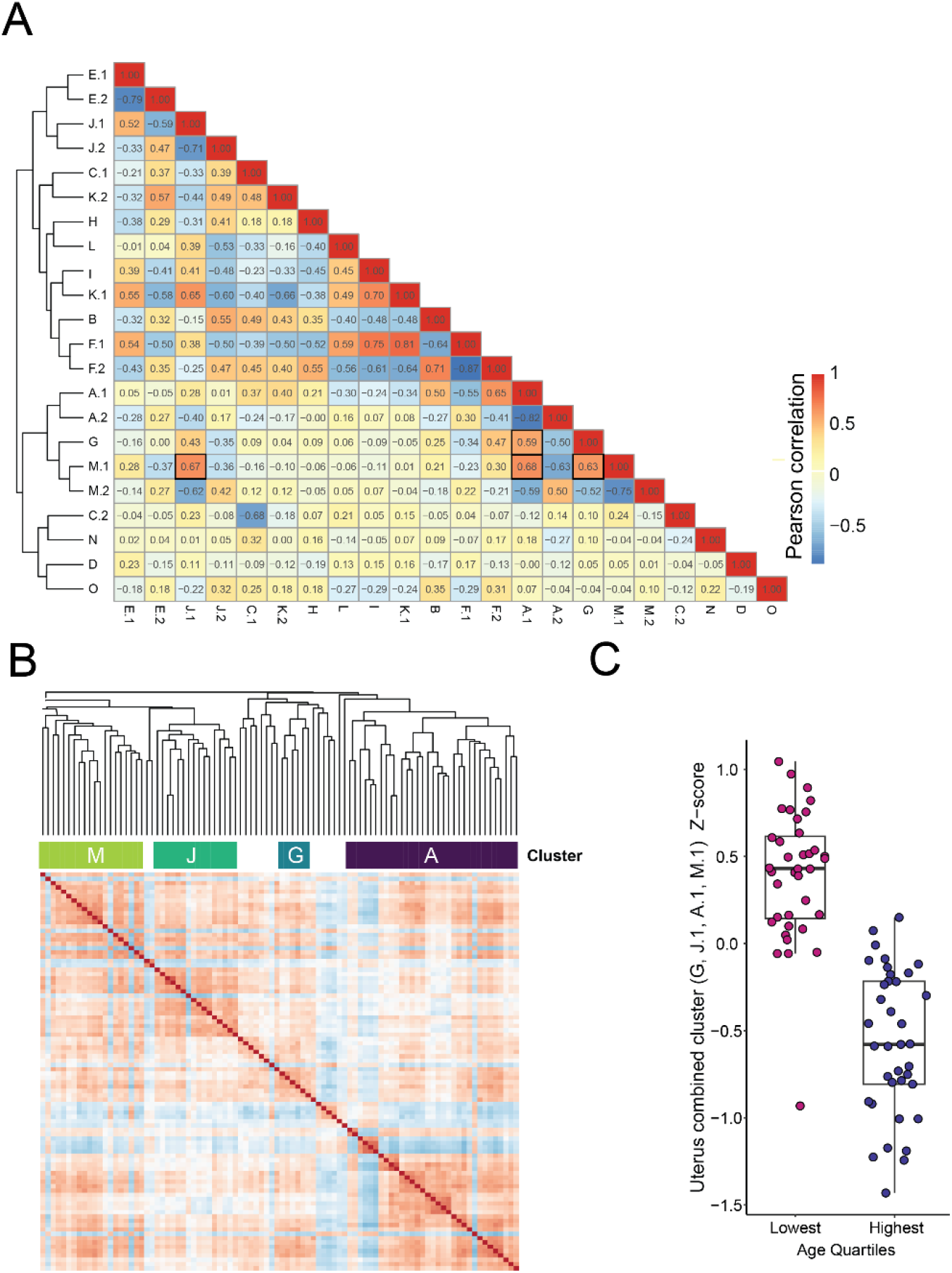
Correlation of uterus clusters and patient age. **(A)** All inter-cluster Pearson correlations in the uterus calculated from cluster Z-scores. The age-related cluster correlations between A1, G, J.1, and M1 are highlighted by black rectangles. **(B)** Kendall correlations from the overall uterus variable gene cluster showing shared clustering across these 4 clusters. (**C)** A combined metacluster of 4 uterus clusters shows strong correlations to patient pre-menopausal age and post-menopausal age (age thresholds are <39.25 and >58 respectively; N = 36, 37).

## Notes

### Competing Interest Statement

The authors have declared no competing interest.

